# Inter-animal variability in activity phase is constrained by synaptic dynamics in an oscillatory network

**DOI:** 10.1101/2022.01.27.478028

**Authors:** Haroon Anwar, Diana Martinez, Dirk Bucher, Farzan Nadim

## Abstract

The levels of voltage-gated and synaptic currents in the same neuron type can vary substantially across individuals. Yet, the phase relationships between neurons in oscillatory circuits are often maintained, even in the face of varying oscillation frequencies. We examined whether synaptic and intrinsic currents are matched to maintain constant activity phases across preparations, using the lateral pyloric (LP) neuron of the stomatogastric ganglion of the crab, *Cancer borealis*. LP produces stable oscillatory bursts upon release from inhibition, with an onset phase that is independent of oscillation frequency. We quantified the parameters that define the shape of the synaptic current inputs across preparations and found no linear correlations with voltage-gated currents. However, several synaptic parameters were correlated with oscillation period and burst onset phase, suggesting they may play a role in phase maintenance. We used the dynamic clamp to apply artificial synaptic inputs and found that those synaptic parameters correlated with phase and period were ineffective in influencing burst onset. Instead, parameters that showed the least variability across preparations had the greatest influence. Thus, parameters that influence circuit phasing are constrained across individuals, while those that have little effect simply co-vary with phase and frequency.

## Introduction

Sensory representations and motor outputs are characterized by the relative timing between different circuit neurons, particularly during oscillatory activity (Ainsworth et al., 2012). Distinct phases of activity within each cycle are found both during oscillations associated with cognition and various behavioral states (Hasselmo et al., 2002; Hajos et al., 2004; Somogyi and Klausberger, 2005; Buzsaki and Wang, 2012; Wilson et al., 2015; Buzsaki and Tingley, 2018; Dragoi, 2020), and during rhythmic motor activity, where they underlie the sequential activation of different groups of muscles (Vidal-Gadea et al., 2011; Bucher et al., 2015; Grillner and El Manira, 2015; Katz, 2016; Kiehn, 2016; Bidaye et al., 2018; Grillner and El Manira, 2020). The relative timing (phase) of a neuron’s activity within each oscillation cycle is dependent on an interplay of intrinsic membrane currents and total cycle-to-cycle synaptic input (Harris-Warrick, 2002; Oren et al., 2006; Marder, 2011; McDonnell and Graham, 2017; Martinez et al., 2019b). There are two confounding aspects of this interplay. First, in many oscillatory systems, phase is maintained over a range of frequencies, i.e., intrinsic and synaptic properties have to ensure that absolute timing of responses changes proportionally to the speed of rhythmic circuit activity (Grillner, 2006; Mullins et al., 2011; Zhang et al., 2014; Le Gal et al., 2017; Martinez et al., 2019b). Second, phase can be very similar across individual animals despite substantial variability in the individual ionic and synaptic currents (Bucher et al., 2005a; Marder and Goaillard, 2006; Calabrese et al., 2011; Marder, 2011; Roffman et al., 2012; Golowasch, 2014; Hamood and Marder, 2014; Marder et al., 2014a; Calabrese et al., 2016).

The phenomenon that circuit activity is maintained despite substantial variability in underlying conductances has been explored most thoroughly in invertebrate central pattern generators, including those of the crustacean stomatogastric ganglion (STG). In these circuits, the timing of neural activity is critically dependent on voltage-gated ion channels (Harris-Warrick et al., 1995b; Harris-Warrick et al., 1995a; Kloppenburg et al., 1999). However, such voltage-gated conductances and the associated ion channel expression show substantial inter-individual variability (Liu et al., 1998; Golowasch et al., 2002; Marder and Goaillard, 2006; Schulz et al., 2006; Marder, 2011; Hamood and Marder, 2014; Marder et al., 2014a), raising the question how activity can be so similar across preparations. A possible explanation is suggested by the finding that voltage-gated conductances do not vary independently, but in a cell type-specific correlated manner (Khorkova and Golowasch, 2007; Schulz et al., 2007; Ransdell et al., 2012; Temporal et al., 2012; Tran et al., 2019). Theoretical work suggests that homeostatic, compensatory tuning explains correlation of expression levels of different ion channels (Prinz et al., 2004a; O’Leary et al., 2013; O’Leary et al., 2014; Franci et al., 2020), and there is some experimental evidence that co-regulation of voltage-gated conductances can have compensatory function to preserve circuit activity (MacLean et al., 2003; MacLean et al., 2005; Ransdell et al., 2012; Zhao and Golowasch, 2012; Ransdell et al., 2013; Santin and Schulz, 2019).

Synaptic currents also vary substantially across individuals and their magnitude is correlated with relative timing of the postsynaptic neuron (Goaillard et al., 2009). In theoretical work, the magnitude of synaptic currents has been varied and tuned alongside voltage-gated conductances to show which combinations and possible mechanisms give rise to similar activity (Prinz et al., 2004a; O’Leary et al., 2014), and it has been suggested that the relative synaptic strengths must be different in individual animals to produce observed activity phases (Gunay et al., 2019). However, it is unknown whether synaptic currents co-vary with individual voltage-gated currents in a correlated manner to compensate for variability in intrinsic neuronal excitability. Furthermore, the effect of synaptic input on rhythmic patterns is not just dependent on synaptic strength, but also on timing, duration, and details of the temporal trajectory of the synaptic current (Prinz et al., 2003; Martinez et al., 2019b).

We examine how synaptic inputs contribute to phase constancy under normal biological conditions in the face of variability across individuals. For this, we use the identified lateral pyloric (LP) neuron in the STG, a follower neuron of the triphasic oscillatory pyloric circuit, which has a single copy in each animal. We examine the variability of synaptic input to the LP neuron across animals and compare that with its activity phase. We examine correlations among synaptic parameters and between these parameters and intrinsic voltage-gated currents of the LP neuron. We then use the dynamic clamp technique to explore how synaptic parameters influence the activity phase of the LP neuron.

## Results

### Variability of phase

The goal of this study was to identify mechanisms that allow a follower pyloric neuron to maintain constant activity phase across preparations, despite considerable variability in cycle period, synaptic input, and voltage-gated conductances. We chose the LP neuron to explore these mechanisms, because it exists as a single copy and is readily identifiable. The LP neuron does not have intrinsic oscillatory activity but receives periodic inhibitory synaptic input from the pacemaker neurons AB and PD, and the follower PY neurons. In each cycle, it rebounds from inhibition to produce a burst of action potentials (Figure 1A).

**Figure 1.**
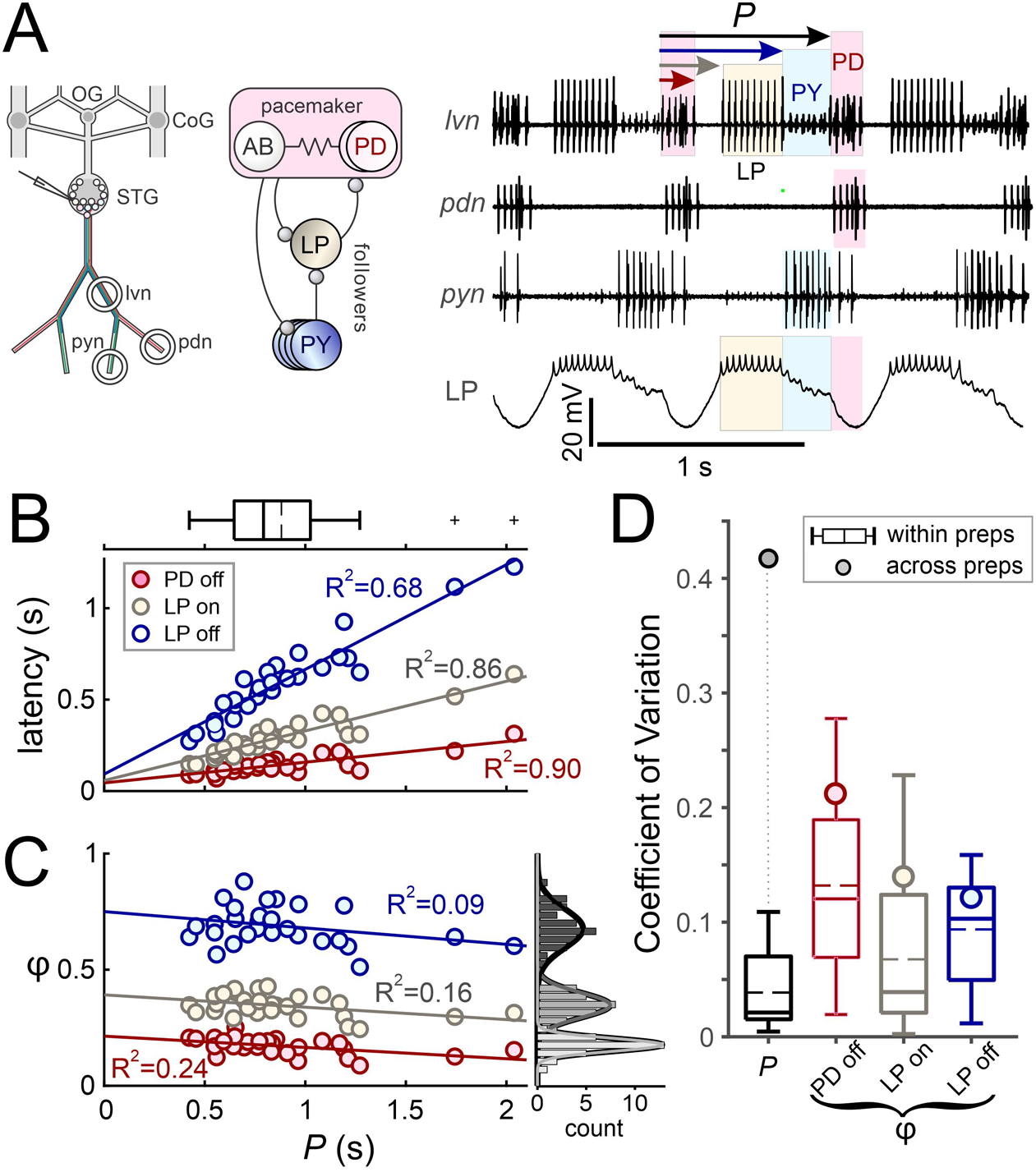
Pyloric activity phases are variable but not correlated with the cycle period. **A**. Left: schematic diagram shows the layout of the STNS in vitro and the locations of intracellular (electrode) and extracellular (circles) recordings. OG: esophageal ganglion, CoG: commissural ganglion, STG: stomatogastric ganglion, lvn: the lateral ventricular nerve, pdn: pyloric dilator nerve, pyn: pyloric nerve. Middle: schematic diagram shows a simplified circuit diagram of the neurons recorded. Ball-and-stick symbols are inhibitory chemical synapses. Resistor denotes electrical coupling. Right: simultaneous extracellular and intracellular recordings show the regular triphasic oscillations of the pyloric circuit. Shown are bursting activity of pacemaker neuron PD and follower neurons LP and PY. Cycle period (*P*) and latencies of the onset and end of each burst (arrows) are calculated from the onset of the PD burst. Extracellular recordings are from the lvn (showing the LP, PD and PY spikes), the pdn (showing the PD spikes) and pyn (showing the PY and LPG spikes). Intracellular recording from the LP neuron shows bursting activity (blue) and slow wave oscillations, as well as timing of IPSPs from the PY (green) and PD (pink) neurons. **B**. Burst latencies of the PD and LP neuron in reference to PD burst onset, as marked in panel A, shown vs. P. Quartile plot shows the distribution of **P**, with the dashed line indicating the mean value. Lines indicate best linear fit, showing that latencies grow proportionally with **P**. **C**. Phase values (φ = latency / *P*) shown vs. *P*. Histograms show distribution of φ values. Linear fits indicate a lack of correlation between all φ values and *P*. *D*. Coefficient of variation of *P* and φ values shown to compare variability of the values within preparations (quartile plots) to their variability across preparations (circles).

The triphasic pyloric activity pattern was continuously present in all preparations, with the temporal sequence of each PD burst being followed after some delay by the LP burst, and then the PY burst (Figure 1A). To quantify the variability in phase and its consistency across different cycle periods (*P*), we measured the latencies of the LP neuron burst onset (LP_on_) and termination (LP_off_) across 28 preparations, from at least 30 s of pyloric activity in each. All latencies were measured with respect to the burst onset of the pacemaker group PD neurons. We also kept track of the burst end phase (PD_off_) of the PD neurons, to quantify the degree to which the pacemakers maintain a constant duty cycle (Abbott et al., 1991). We did not quantify the PY neuron burst onset and end phases, because in *C. borealis*, they are virtually identical to LP_off_ of the same cycle, and PD_on_ of the subsequent cycle (Goaillard et al., 2009). First, we determined the mean values for latencies, *P*, and phases (*φ* = latency/*P*) in each preparation. Across preparations, *P* ranged from 423 to 2038 ms, with a mean of 880 ms (± 368 SD). As reported previously (Bucher et al., 2005a; Goaillard et al., 2009), the latency values of PD_off_, LP_on_ and LP_off_ increased roughly proportionally with *P* (Figure 1B). Consequently, phases did not change significantly with *P* (Figure 1C).

It is noteworthy that a lack of correlation with *P* does not mean that phases were completely invariant, as the histograms in Figure 1C indicate. We compared the variability of mean phases and *P* across preparations with the cycle-to-cycle variability observed across individual preparations. Figure 1D shows box plots of coefficients of variation (CVs) within individual preparations, alongside the single CV values calculated from the means across preparations. Variations in phase were in the same range within and across preparations. In contrast, there was a much larger variability of mean *P* across preparations than within each preparation. These results confirm that phases are under much tighter control across preparations than cycle period.

### Inter-individual variability of voltage-gated currents and synaptic inputs

The maximal conductances (*g*_max_) of voltage-gated ionic (henceforth called intrinsic) currents in identified pyloric neurons, including LP, show large variability across animals (Marder and Goaillard, 2006; Schulz et al., 2006; Goaillard et al., 2009; Marder, 2011; Golowasch, 2014; Marder et al., 2014a). Variability of *g*_max_ is well correlated with variability in transcript levels of the underlying ion channel genes, and therefore serves as a good proxy for variability of ion channel numbers (Schulz et al., 2006).

We measured intrinsic currents in LP for two reasons. First, variability has previously only been determined for *g*_max_, and we wanted to also examine the variability of voltage-dependence. Second, we measured synaptic currents in the same preparations to establish if there was co-variation that could indicate compensatory regulation of intrinsic and synaptic currents. We performed these measurements of synaptic and intrinsic currents during ongoing rhythmic pyloric activity, restricting ourselves to the subset of intrinsic currents that under these conditions can be measured without pharmacological manipulation (Zhao and Golowasch, 2012). They included the high-threshold voltage-gated K^+^ current (*I*_HTK_), the transient K^+^ current (*I*_A_), and the hyperpolarization-activated inward current (*I*_H_) (Figure 2A). Currents were converted to conductance values, and for K^+^ currents, the activation curves in each individual preparation were fit with a sigmoid to determine *g*_max_, voltage of half activation (*V*_1/2_), and the slope factor (*k*) (Figure 2B). For *I*_HTK_, we obtained these parameters for both the transient (*I*_HTKt_) and the persistent (*I*_HTKp_) components.

**Figure 2.**
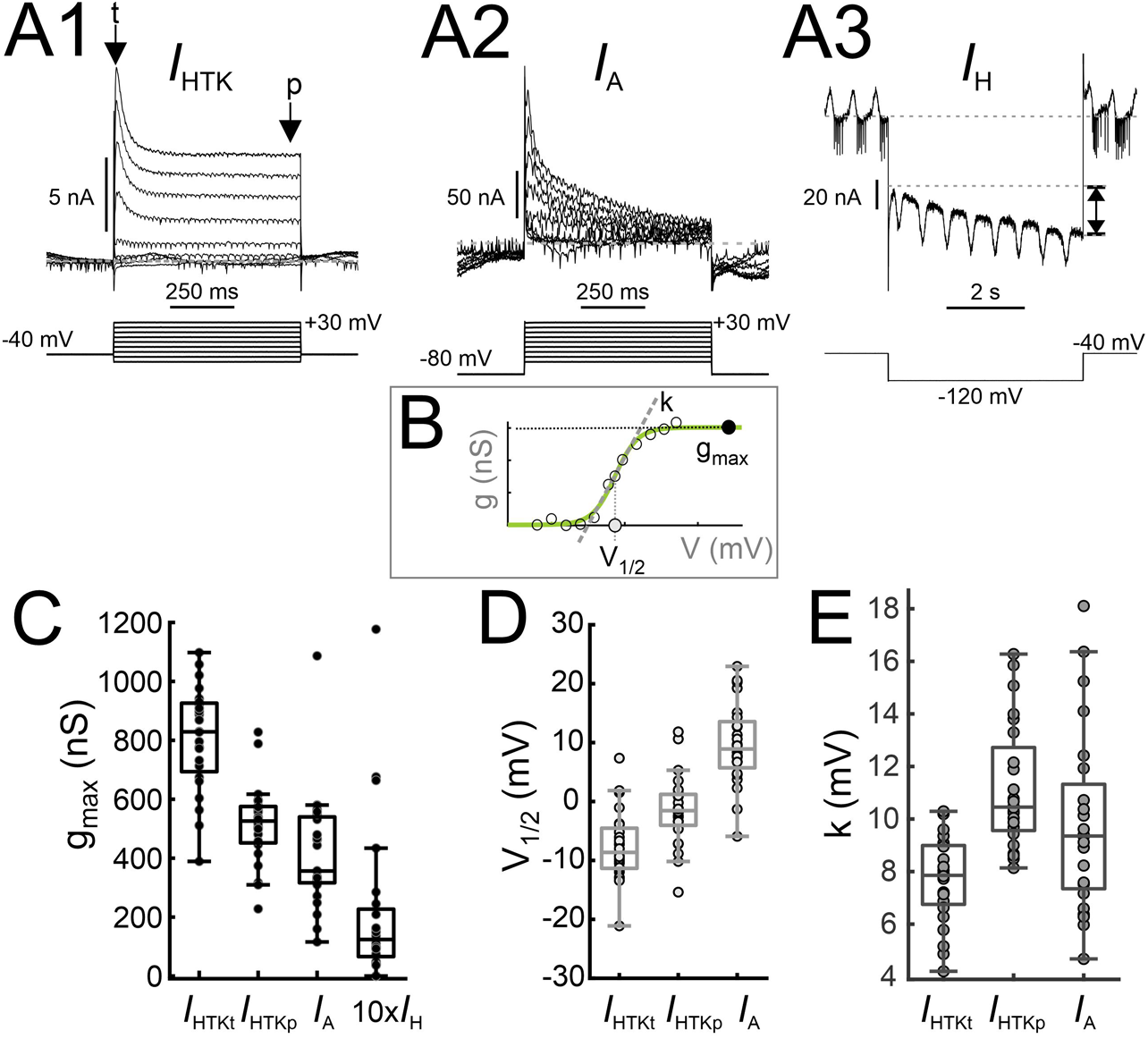
Parameters defining voltage-gated currents *I*_HTK_, *I*_A_ and *I*_H_ show considerable variability. **A.** Example voltage clamp recordings of high-threshold potassium currents (*I*_HTK_, **A1**; arrows indicate the transient [t] and persistent [p] components), the transient potassium A current (*I*_A_, **A2**) and the H current (*I*_H_, **A3**). Double-arrow in A3 indicates the measured amplitude of *I*_H_. **B.** Schematic diagram of fits in each experiment to the *I*_HTK_ and *I*_A_ conductances (g, measured by dividing current by the driving force, assuming *E*_K_ = −80 mV). Fits were used to calculate maximum conductance (g_max_), half-activation voltage (V_1/2_) and activation slope factor (k, measured from the slope of the dashed line). **C.** Maximal conductances of the transient and persistent components of *I*_HTK_, *I*_A_ and *I*_H_ across preparations. **D.** Half activation voltage of the transient and persistent components of *I*_HTK_ and of *I*_A_ across preparations. **E.** Activation slope factor of the transient and persistent components of *I*_HTK_ and of *I*_A_ across preparations.

Like previous reports (Schulz et al., 2006; Khorkova and Golowasch, 2007), *g*_max_ values of *I*_HTK_, *I*_A_, and *I*_H_ showed large variability (Figure 2C). In addition, we found that for both *I*_HTK_ and *I*_A_, the parameters *V*_1/2_ and *k* were also subject to large variability (Figure 2D-E). We interpret this as an indication that not only the number of channels, but also their gating properties can vary substantially across individuals.

To examine variability of synaptic inputs across preparations, we recorded the LP neuron’s graded inhibitory postsynaptic currents (IPSCs) in response to PD and PY neuron input during ongoing pyloric activity (Figure 3A). The shape of the recorded IPSCs varied considerably across preparations. We used 12 parameters to quantify the IPSC characteristics (Figure 3B; see Methods). The distributions of these parameters showed that the IPSC in the LP neuron varies greatly across preparations (Figure 3C and Table S1).

**Figure 3.**
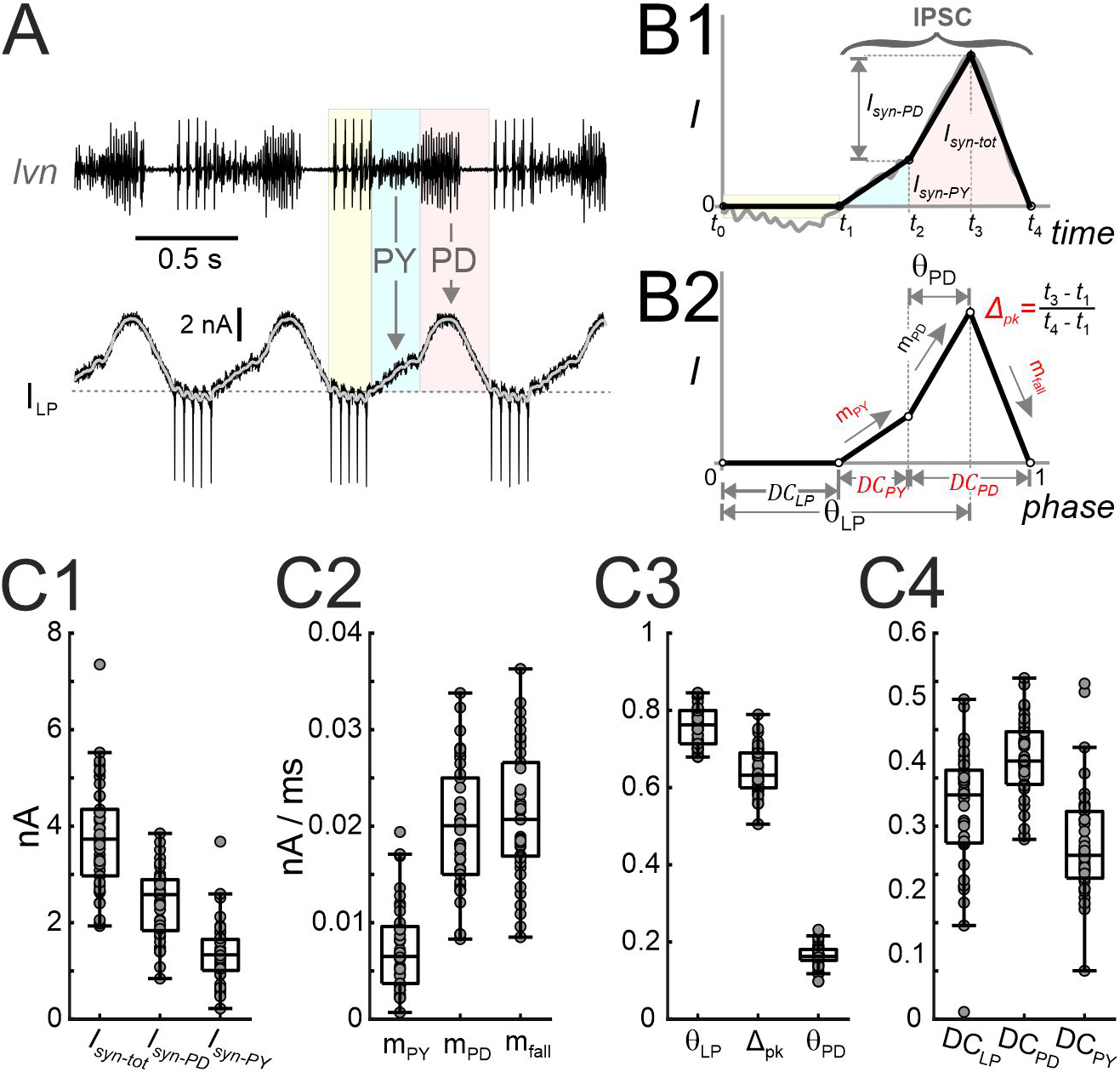
Parameters that define the synaptic input show considerable variability. **A**. Total current measured in the LP neuron voltage clamped at a holding potential of −50 mV during the ongoing pyloric rhythm. The pyloric rhythm is recorded extracellularly (*lvn*), indicating the timing of the LP, PY and PD neuron bursts. LP action potentials escape the voltage clamp and can be seen in the current and extracellular recordings (pale yellow). The portion of the current outside this range is due to synaptic input (downward arrows) from the PY (light blue) and pacemaker (PD, pink) neurons. The gray curve is the current low-pass filtered (<20 Hz). **B1.** The synaptic waveform shape (gray curve) during a single cycle of oscillation is approximated by a piecewise-linear curve (black curve), marked by five time points (t_0_-t_4_) denoting the borders of the colored regions in panel A. The time range of the IPSC and the amplitudes of the synaptic currents due to the pacemaker neurons (*I*_*syn-PD*_), due to the PY neurons (*I*_*syn-PY*_), and the sum of the two (*I*_*syn-tot*_) are marked. **B2.** The piecewise-linear curve of B1 shown in phase (time/period). This normalized curve is used to define the parameters of synaptic input to the LP neuron. For definitions, please refer to the main text. Five primary parameters (in red) are chosen for further analysis. C. The inter-individual variability of different synaptic parameters, including current amplitudes (**C1**), slopes (**C2**), peak phases (**C3**) and duty cycles (**C4**).

The latency of the LP burst onset relative to the pacemakers is shaped by the interaction between its intrinsic voltage-gated ionic currents and the synaptic input that it receives. Notably, hyperpolarization during inhibition de-inactivates *I*_A_ and activates *I*_H_ (Harris-Warrick et al., 1995b; Harris-Warrick et al., 1995a). This plays an important role in controlling the timing of the burst onset, because *I*_H_ increases the strength of the rebound burst and advances its onset, while *I*_A_ delays it (MacLean et al., 2005). The activation levels of *I*_H_ and *I*_A_ in each cycle depend on the strength, duration, and history of the inhibition. In addition, *φ*_LP on_ is sensitive to changes in both magnitude and temporal trajectory of synaptic inputs (Goaillard et al., 2009; Martinez et al., 2019b). We hypothesized that the synaptic inputs to the LP neuron may covary in a compensatory fashion with its intrinsic properties, thus resulting in a relatively constrained activity phase across animals. We therefore examined the extent to which the synaptic input parameters may be coregulated with *g*_max_ of these ionic currents, as well as *I*_HTK_. We also tested for any correlations of synaptic parameters with *V*_1/2_ and *k* values of the K^+^ currents. We did not find any significant pairwise linear correlations between any of the synaptic and intrinsic current parameters (Figure 4 and Table S2; all P values in linear regression analysis > 0.05; N = 19).

**Figure 4.**
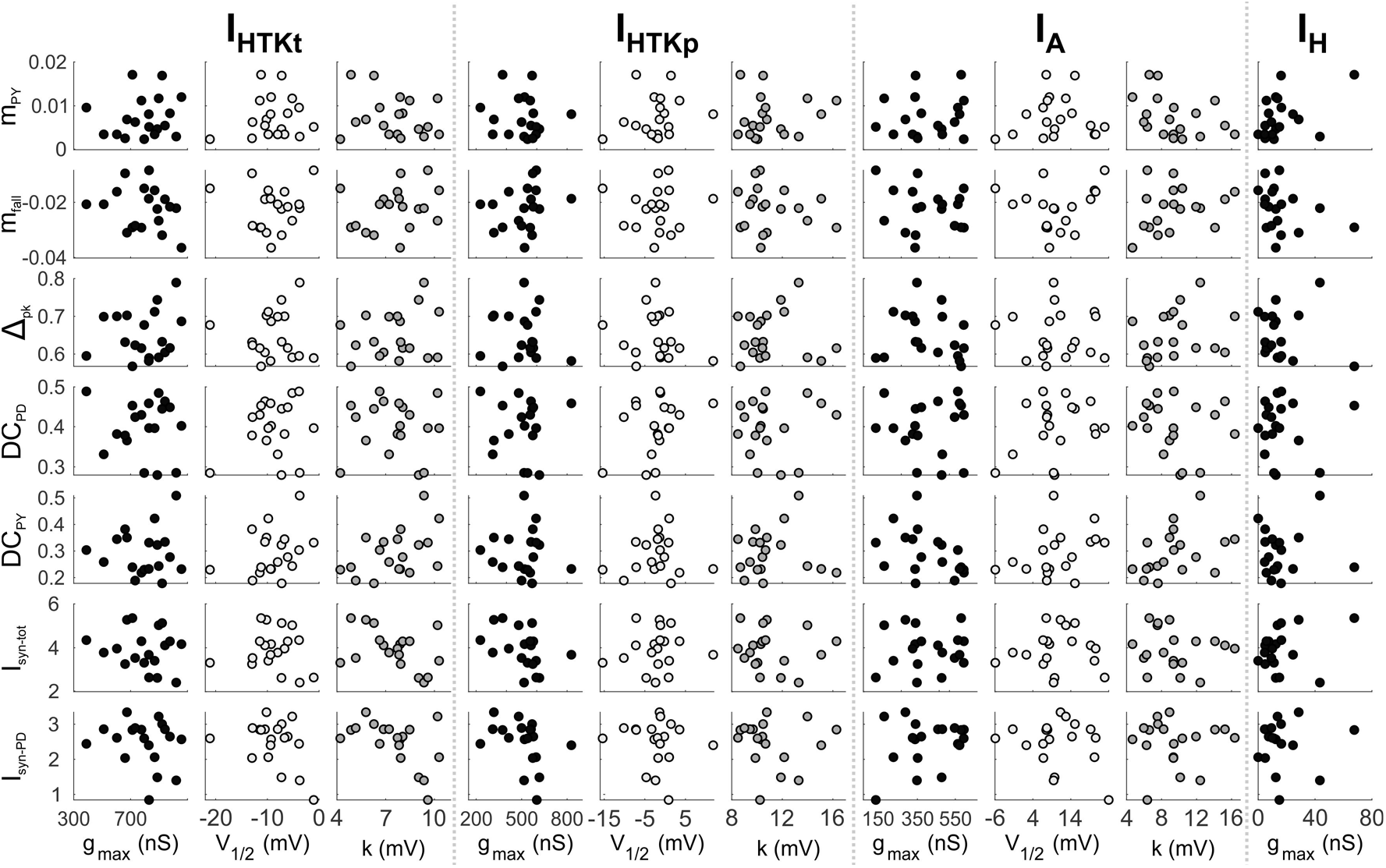
There are no pairwise linear correlations between any of the synaptic parameters and parameters of the voltage-gated ionic currents.

### The LP burst onset is influenced by synaptic parameters

Our results suggest that the consistency of phase across individuals and its independence of cycle period do not simply arise from pairwise correlations between synaptic and intrinsic parameters. We therefore asked if individual synaptic or intrinsic current parameters are good candidates for playing a substantial role in controlling phase. To this end, we made use of the variability of mean *P* and the limited variability of mean *φ*_LP on_ across individuals and performed correlational analyses. For synaptic currents, we included the maximum IPSC amplitude and the amplitude of the pacemaker IPSC, as these are commonly used synaptic parameters. Otherwise, we restricted the analysis to the non-redundant set of parameters. As described in the Methods, the 5 non-redundant parameters are the subset of measures that are sufficient to describe synaptic current trajectory and can theoretically vary independently of each other.

First, we tested whether variability of current parameters was correlated with *P*. We found that a subset of the parameters describing the trajectory of synaptic currents, but none of the intrinsic parameters, showed correlations with *P* (Figure 5). The IPSC slope parameters *m*_PY_ and *m*_fall_ were strongly correlated with *P*, but in opposite directions. *DC*_PD_ also showed a weak (negative) correlation with *P*. The latter is somewhat surprising, as we found a negative trend but no correlation between *φ*_PD off_ and *P* in the pyloric pattern analysis shown in Figure 1C.

**Figure 5.**
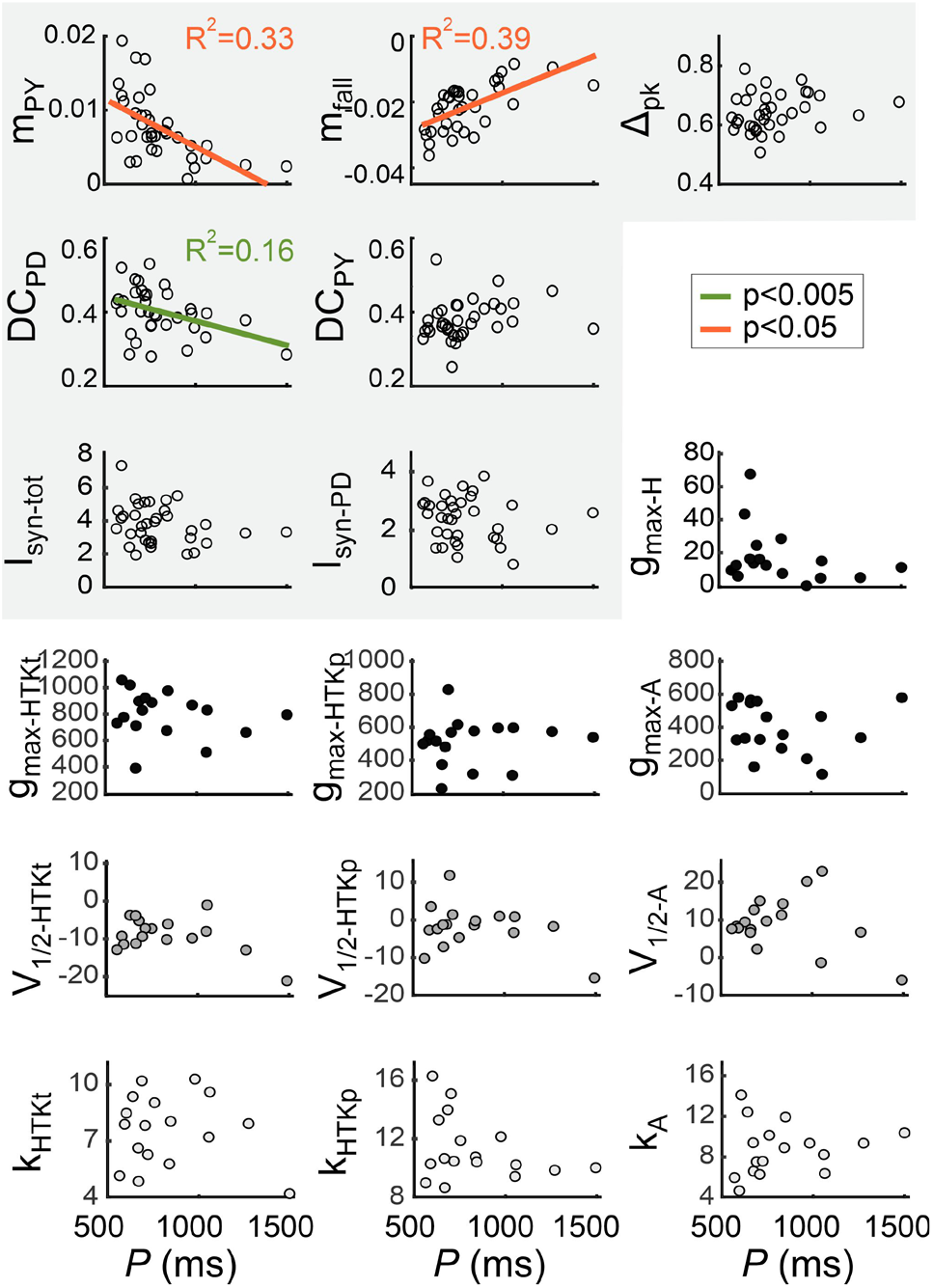
A subset of the LP neuron synaptic, but not intrinsic, parameters are correlated with the pyloric cycle period. The five primary synaptic parameters, the amplitudes of the total and pacemaker-component of the synaptic current, and the intrinsic current parameters are compared with the pyloric cycle period (*P*) across preparations. Three synaptic parameters, but no intrinsic parameter, covary with *P*. The synaptic parameters are highlighted.

Next, we explored whether *φ*_LP on_ was correlated with any of the current parameters (Figure 6). Once again, we found correlations with a subset of the parameters describing the trajectory of synaptic currents, but none with intrinsic parameters. *φ*_LP on_ was weakly positively correlated with *m*_PY_, and strongly negatively correlated with *Δ*_pk_. Interestingly, *φ*_LP on_ was correlated strongly with both *DC*_PD_ and *DC*_PY_, with opposite signs. This suggests that synaptic inputs from both the pacemakers and the PY neurons may influence *φ*_LP on_, even though the input from the PY neurons is primarily responsible for the termination of the LP neuron burst, not its onset (Marder and Bucher, 2007). In comparison with Figure 5, *m*_PY_ and *DC*_PD_ were correlated with both *φ*_LP on_ and *P*, whereas *m*_fall_ was only correlated with *P*, and *Δ*_pk_ and *DC*_PY_ only with *φ*_LP on_.

**Figure 6.**
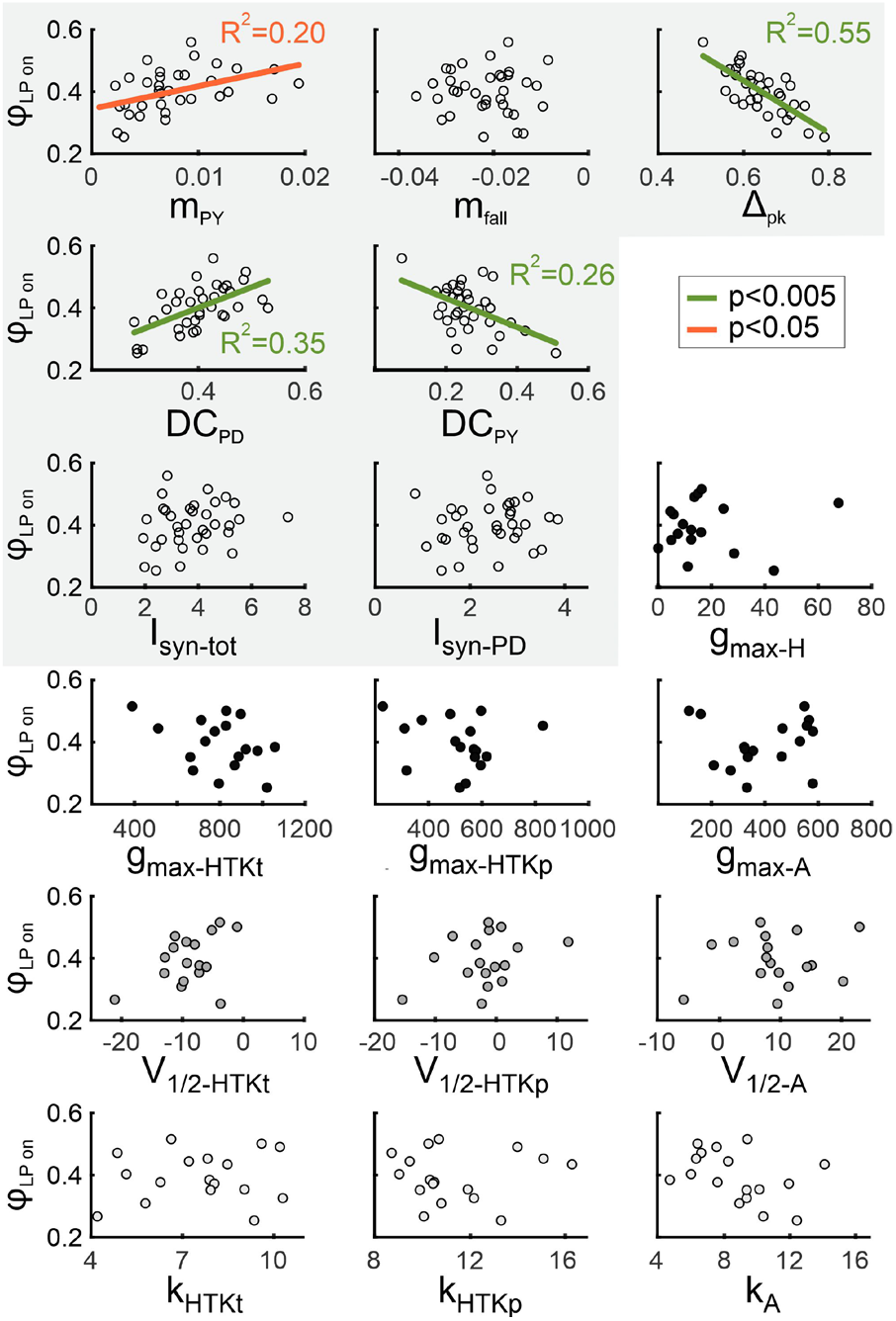
The LP neuron burst onset phase is correlated with multiple synaptic, but not intrinsic, parameters. The LP burst onset phase (*φ*_LP on_) is compared with the five primary synaptic parameters, the amplitudes of the total and pacemaker-component of the synaptic current, and the intrinsic current parameters across preparations. *φ*_LP on_ covaries with four synaptic parameters, but not with intrinsic parameters. The synaptic parameters are highlighted.

### The influence of synaptic parameter variation on the LP neuron’s burst onset

Given that our results revealed no correlations between single intrinsic current parameters and *φ*_LP on_, but correlations between *φ*_LP on_ and several synaptic parameters, we further explored which aspects of the overall synaptic current trajectory were important. Because we found no correlations between synaptic current amplitudes and *φ*_LP on_ or *P*, we restricted the analysis to the non-redundant parameters. As stated above, these 5 parameters can theoretically be varied independently to change synaptic current trajectory. However, this does not mean that they actually varied independently in the measured experimental data. Indeed, we found that most parameter pairs were correlated, some strongly and others weakly (Figure 7A). In particular, *Δ*_pk_ and *DC*_PD_ were strongly correlated, as was expected for parameters that quantify the contribution of the pacemakers. However, *m*_fall_, which also depends on the strength and the timing of the pacemaker inputs, was not correlated with *Δ*_pk_ or *DC*_PD_. Surprisingly though, *m*_fall_ was strongly correlated with *m*_PY_ and, consistent with this fact, *Δ*_pk_ and *DC*_PD_ were also correlated with *m*_PY_.

**Figure 7.**
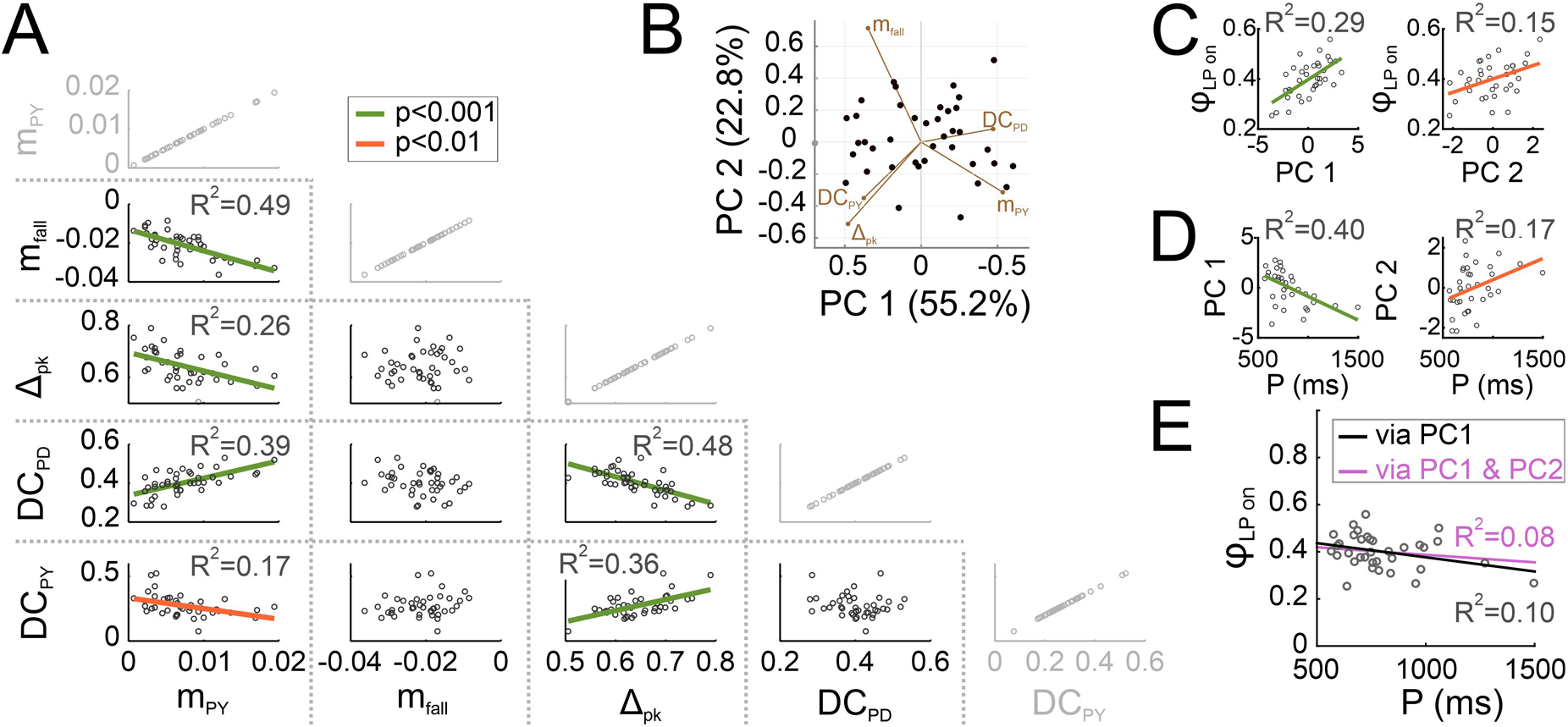
The primary synaptic parameters are correlated. **A.** The five primary synaptic parameters were compared pairwise across preparations. Of the 10 non-trivial comparisons (shown in black), 6 showed significant correlations. The trivial comparisons (gray) are shown for clarity. **B.** Principal component analysis was used to find directions of largest variability among the five synaptic parameters. The first two principal components described 78% of the variability in synaptic parameters. Filled circles show all recorded synaptic waveforms, projected down to the PC1-PC2 plane. Percentages on axis labels indicate the extent of variability in the direction of the PC. The directions of the five primary synaptic parameters in the PC1-PC2 plane are indicated by brown line segments. C. Across preparations, the LP burst onset phase (φ_LP_) is correlated with both PC1 and PC2 (but not PC3, PC4 or PC5). **D**. Across preparations, both PC1 and PC2 (but not PC3, PC4 or PC5) are correlated with the pyloric cycle period (*P*). **E.** Using the PC1 and PC2 correlations with φ_LP_ and *P* (lines in left graphs of panels C and D) to calculate a linear relationship (black line) between φ_LP_ and *P* correctly predicts a lack of correlation between these two factors. Including both the PC1 and PC2 correlations (all lines in panels C and D) to do the linear prediction (magenta line) does not greatly improve the prediction.

Our correlational analysis indicates that multiple synaptic parameters co-vary across experiments. The correlation of multiple synaptic parameters with *φ*_LP on_ (as seen in Figure 6) indicates that these five parameters in fact covary across preparations and therefore the variation in synaptic shape occurs in a lower dimensional parameter space. To address this issue, it is possible to simplify the correlational analysis by determining which combination of parameters explains the observed variability in *φ*_LP on_. To reduce the dimensionality of the IPSC parameter space, we performed principal component analysis (PCA). We found that 95% of the total variability of IPSC parameters were explained by the first three PCs (PC1: 55.2%; PC2: 22.8%; PC3: 16.5%). Figure 7B shows all synaptic waveforms in the plane of the first two PCs, as only PC1 and PC2 were significantly correlated with *φ*_LP on_ (PC1: p < 0.001; PC2: p =0.013; Figure 7C). Interestingly, both PC1 and PC2 (and only these) were also significantly correlated with *P* (PC1: p < 0.001; PC2: p =0.024; Figure 7D).

To examine whether the coordinated variation of synaptic parameters in the direction of PC1 was sufficient to explain phase maintenance, we used the linear regression fit equations of PC1 vs. *φ*_LP on_ and PC1 vs. *P* (left panels of Figure 7C-D) to predict a linear relationship between *φ*_LP on_ and *P*. In Figure 7E, we compare this prediction (black line) with the data for LP on over P shown in Figure 1C (open circles). This comparison produced a coefficient of determination of R^2^ = 0.10, which was comparable with the linear fit obtained in Figure 1C (R^2^ = 0.16 for *φ*_LP on_). This indicates that variation of the synaptic conductance trajectory with *P* along PC1 is sufficient to remove the correlation between *φ*_LP on_ and *P*, thus predicting phase maintenance across preparations. Additionally correcting this prediction by adding the linear regression fit equations of PC2 (right panels of Figure 7C-D) did not greatly change this prediction (violet line in Figure 7E, R^2^ = 0.08).

Our analysis of data obtained during spontaneous pyloric rhythmic activity revealed combinations of synaptic parameters whose coordinated variation could potentially result in relatively constant *φ*_LP on_ across preparations, despite variation in *P*. However, there are two caveats. First, correlation may result from causation in some cases, but not in others. A synaptic parameter (or a principal component, such as PC1) that is correlated with *φ*_LP on_ may in fact causally influence *φ*_LP on_. If so, the system must adjust this parameter at different cycle periods in order to produce phase maintenance. For example, this could explain why PC1 is correlated with both *φ*_LP on_ and *P*. In contrast, a parameter may simply change with *φ*_LP on_ but not influence it, in which case its change with *P* would not contribute to phase maintenance. Similarly, a parameter such as *Δ*_pk_ that is correlated with *φ*_LP on_ but not *P* may also causally influence *φ*_LP on_ (see, e.g., Martinez et al., 2019b) and would therefore be kept constant across animals in order to maintain phase. Second, causation may not necessarily reveal itself as a correlation. In our data, *φ*_LP on_ (and all other pyloric phases) varied in a fairly limited range, independent of the large variability of *P*. Therefore, simply analyzing correlations in data obtained from spontaneous rhythms constrained the maximum effect a synaptic parameter may have on *φ*_LP on_ to the same limits. Within these limits, a parameter that has no correlation with *P* or *φ*_LP on_ may in fact have a strong influence on *φ*_LP on_ and, for this reason, be kept constant across animals (and thus show no correlation with *P*).

For these reasons, establishing a causal influence of synaptic parameters on *φ*_LP on_ requires experimentally controlling and systematically varying them. To this end, we performed a set of experiments in which the LP neuron was synaptically isolated, and synaptic conductance waveforms were artificially applied using the dynamic clamp technique. Conductance trajectories were constructed to resemble the current trajectories and adhering to the same decomposition into parameters shown in Figure 3B. We kept the cycle period constant at 1 s and injected the waveforms periodically until the LP burst activity attained a steady state (~ 30 cycles; Figure 8A). In each experiment, this procedure was repeated with 80 different synaptic trajectories in randomized order (Figure 8B). Because our focus here is on variability and activity phase, we did not do a complete analysis of these dynamic clamp experiments on LP activity and only considered the effect on the LP neuron’s burst onset at steady state, measured as the latency from the end of the artificial synaptic input (inset of Figure 8A; also see Methods).

**Figure 8.**
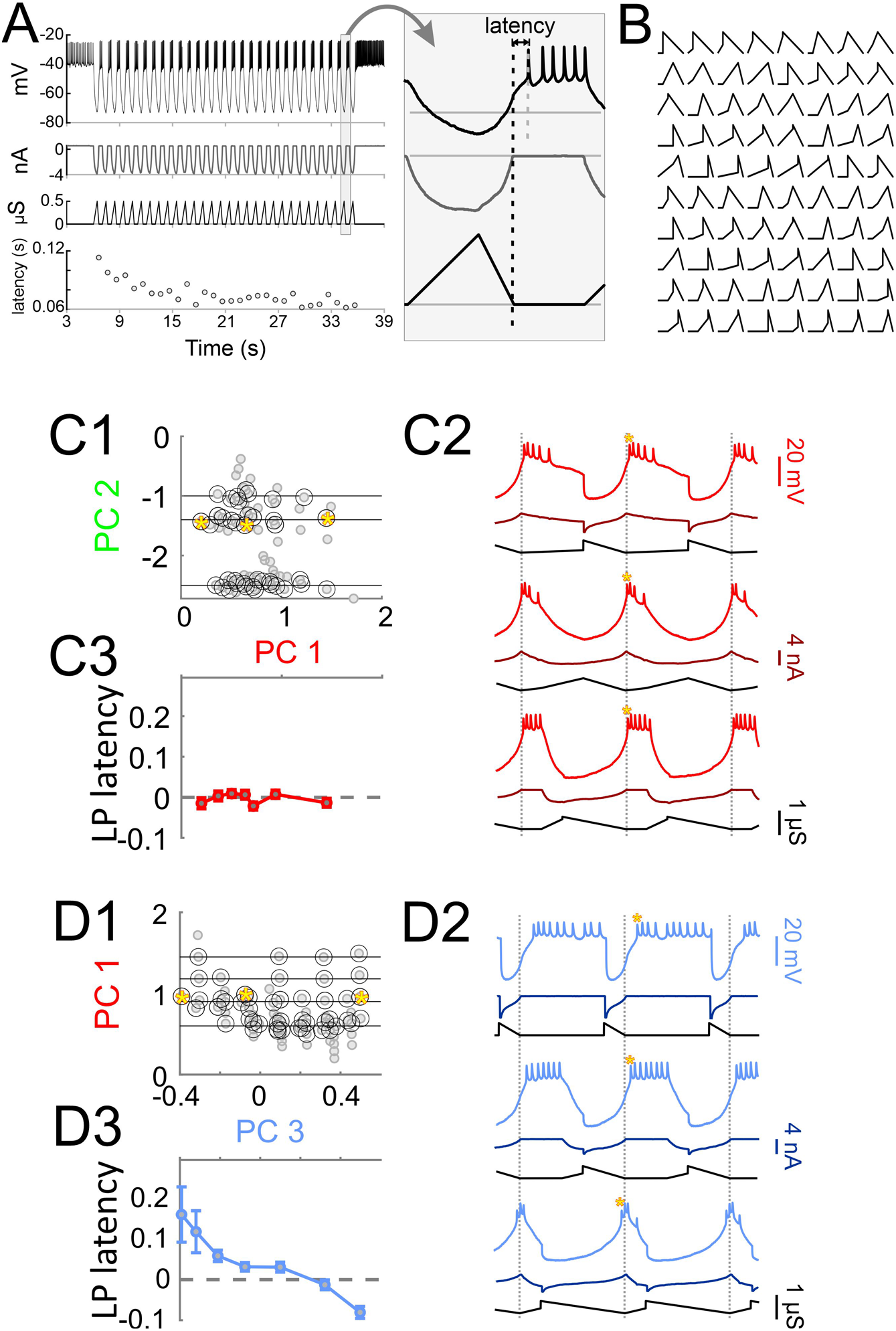
Using dynamic clamp to inject a periodic synaptic conductance waveform into the synaptically-isolated LP neuron to measure the latency of LP burst onset. **A.** A pre-determined conductance waveform (one of 80) is injected into the synaptically-isolated LP neuron as an inhibitory synapse for 30 cycles at a cycle period of 1 s. The latency of the LP burst onset, measured form the end of the conductance waveform (long vertical dashed line in inset), reaches a steady state value after several cycles. Inset shows the last cycle. **B.** 80 synaptic conductance waveforms were used periodic dynamic clamp injection in each LP neuron, as described in A. **C.** An example analysis of the sensitivity of LP burst onset latency to changes in synaptic waveform shape along PC3, while PC1 remains constant (but other PCs vary freely). **C1.** The 80 synaptic waveforms (black circles) used in dynamic clamp experiments were sorted by projecting the shape down to the PC3-PC1 plane. Pairs of waveforms (red circles) that fell along horizontal lines of constant PC1 (and were apart by at least 0.05 in PC3 units) were chosen for sensitivity analysis. Inset above the figure shows the waveform shapes for the gray filled circles. **C2.** Example responses of the LP neuron to dynamic clamp injection of synaptic conductance waveforms marked by the yellow stars in A1. C3. The change in LP burst latency (see A) as a function of the change in PC3 value (in bins of 0.1), averaged across constant PC1. The slope of this change (Δlatency/ ΔPC) is used as a measure of sensitivity. **D.** Same as C, but changing the waveform along PC1 while keeping PC2 constant.

The correlations obtained from the principal component analysis imply that varying synaptic waveform along PC1 while retaining the respective correlations with *P* and *φ*_LP on_ shown in Figure 7C-D should keep *φ*_LP on_ independent of *P*, which would be sufficient to describe phase constancy across preparations. We used our dynamic clamp data to examine whether changing the synaptic waveform along PC1 in fact influenced the LP burst onset latency. To do so, we first described our 80 synaptic waveforms in terms of PC1-PC5. Because visualization of 5D space is difficult, if not impossible, we show the waveform shapes projected down to the PC1-PC2 and PC1-PC3 planes (Figures 8C1 and 8D1, respectively). To analyze the effect of changing the synaptic shape in the direction of each PC, we first measured the sensitivity of the LP burst onset latency to changing that PC, while keeping another PC constant (see Methods). We did this analysis for each pair of PCs. Two examples are shown in Figures 8C and 8D. Surprisingly, the LP burst onset latency showed little sensitivity when the synaptic waveform was changed along PC1 while keeping PC2 constant (example in Figure 8C2; average effects in Figure 8C3; 1-way ANOVA: p=0.26 and F=1.32). In contrast, changing the synaptic waveform along PC3 while keeping PC1 constant produced a very large decrease the burst onset latency (example in Figure 8D2; averages in Figure 8D3; 1-way ANOVA: p<0.001 and F=8.21 using). This result is surprising because it implies that changing the synaptic waveform along PC1 does not result in any change in the LP burst onset, which contradicts our initial interpretation of the correlations observed in Figure 7C-D. If changing the synaptic waveform along PC1 does not produce any change in the LP burst onset, then it makes no sense to claim that the mechanism for phase constancy across preparations with different cycle periods is by changing the synaptic waveform along PC1. Similarly, the synaptic waveforms showed no correlation between PC3 and either *P* or *φ*_LP on_. Yet, experimentally changing the synaptic waveform along PC3 produces a large effect on the LP burst onset.

In Figure 9, we summarize the statistics of the effect of changing the synaptic waveform (with dynamic clamp) along each PC, while keeping one other PC constant. Figure 9A is an illustration of how synaptic waveform changes along each of the five PCs. Figure 9B shows the sensitivity of the LP neuron’s burst onset latency to these changes, either grouped by the PC that was systematically varied while one other was fixed (Figure 9B1) or grouped by the PC that was fixed while one other was systematically varied (Figure 9B2). On average, changing the synaptic waveform along each of the PCs, except for PC2, had some effect on the burst onset latency (Figure 9C). PC3 had the largest effect, followed by PC5. In addition, fixing PC3 made the LP neuron’s burst onset latency insensitive to varying any of the other PCs, while fixing any of the other PCs did not have that effect (Figure 9D). These results suggest that the synaptic parameters (PC1 and PC2) that show the largest variation across preparations have little influence on the burst onset of the LP neuron, whereas two of the synaptic parameters (PC3 and PC5) which have large effect on the burst onset show little variability and are kept relatively constant across preparations.

**Figure 9.**
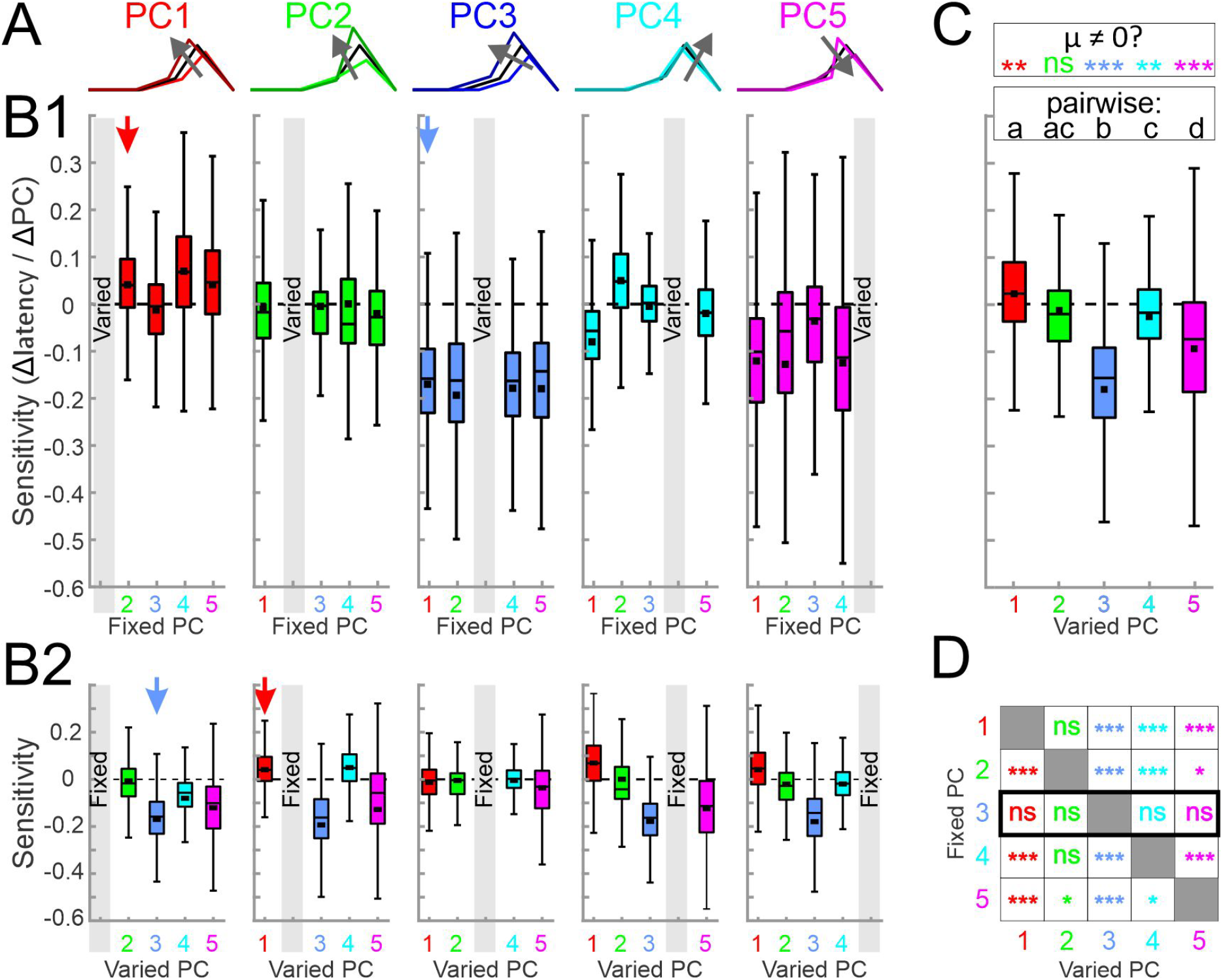
Varying synaptic waveform shape along each principal component while keeping another principal component constant. **A.** Graphical representation of how the shape of a representative synaptic waveform (black) changes when synaptic parameters are shifted by 25% in the direction of each principal component (in the direction of the arrow). **B1**. Sensitivity of LP burst onset latency to changing the principal component along a single PC (marked by gray box in each panel) while a single other PC is kept constant (and the other three are not controlled). Examples of the process (marked by arrows) are shown in Fig. 8C & 8D. In these panels, quartile plots are from data including every individual sensitivity value (200-500 data points) in each experiment (N=10 animals). Black squares show mean values. **B2.** The same data as in B1, reorganized so that each panel shows data when a single PC (gray box) is kept constant while a single other PC is varied (and the other 3 are not controlled). The red and blue arrows point to the same data as they do in B1. **C.** Overall sensitivity of LP burst latency (see Fig. 8) to changing the synaptic waveform along each PC, measured as an overall average of the values shown in panel B. In this graph only the mean sensitivity values in each experiment are used as data points, so that each quartile plot correspond N=10 data points. These sensitivities were significantly different (1-W RM-ANOVA p < 0.001). Different letters (a-d) indicate p<0.01 with post-hoc Tukey test; shared letters indicate p>0.05. Asterisks indicate post-hoc analysis indicating whether the mean value (μ) ≠ 0, **p<0.001, ***p<0.0001. **D.** Statistical summary of panel A data, indicating how varying one PC, while keeping another PC constant (and not controlling others), would produce a change in LP burst latency. Asterisks indicate post-hoc analysis indicating whether μ ≠ 0, *p<0.01, **p<0.001, ***p<0.0001.

## Discussion

### Variability of activity phases within and across individuals

During oscillatory circuit activity, differences in sensory, descending, and modulatory inputs often result in different activity phases between different neurons, whereas similar behavioral settings and circuit states produce characteristic phase relationships (Marder and Bucher, 2001; Wang, 2010; Grillner and El Manira, 2015; Wilson et al., 2015; Frigon, 2017; Grillner and El Manira, 2020). These activity phases can even be maintained over a wide range of rhythm frequencies within individuals, which has been demonstrated in many motor systems (DiCaprio et al., 1997; Wenning et al., 2004; Marder et al., 2005; Grillner, 2006; Le Gal et al., 2017).

Across individuals, activity patterns can vary, particularly in cycle period, but retain enough consistency in the activity phases to be readily matched across the same individuals. In the pyloric circuit, spontaneous *in vitro* rhythmic patterns in individual preparations show some cycle-to-cycle variability in the bursting neurons’ activity phases, consistent with cycle-to-cycle variability in cycle period (Bucher et al., 2005a; Elices et al., 2019). However, mean phases are well maintained when mean cycle period is experimentally altered (Hooper, 1997; Tang et al., 2012; Soofi et al., 2014). Across individuals, phases also show some limited variability but are insensitive to substantial differences in mean cycle period (Bucher et al., 2005a; Goaillard et al., 2009). We confirm here that phases can vary across individuals but do not correlate with mean cycle period (Figure 1C). We also show that the variability of neuronal activity phases across individuals is within the same ranges as cycle-to-cycle variability within individuals, even though cycle period varies substantially more across individuals than it does within individuals (Figure 1D). This raises the question of whether these activity phases are constrained to a small range of variability. We assert that there is no absolute measure for how much variability constitutes a lot or a little, and such an assessment should depend on a reference value. For example, in the leech heartbeat system, variability in phase has been interpreted as being large because phase values varied as a substantial fraction of the reference cycle (Wenning et al., 2018). In the pyloric circuit, variability of phases under control conditions is limited in the sense that phases are largely constrained to values that differ from those under different neuromodulatory conditions (Marder and Bucher, 2007; Harris-Warrick, 2011). In our dataset, variability in phase was large enough to allow us to search for correlations with intrinsic and synaptic current parameters, but the fact that phases remain independent of cycle period justified asking which parameters may be constrained or may co-vary in a compensatory manner in order to achieve consistent circuit output phases across individuals.

### Variability of intrinsic and synaptic currents

Activity phases are shaped by both intrinsic and synaptic currents, both of which can vary substantially across individuals. In the LP neuron, the known intrinsic currents vary several-fold across animals (Liu et al., 1998; Schulz et al., 2006; Schulz et al., 2007; Golowasch, 2014). We found that the voltage-gated currents do not just vary in magnitude, but that the half-activation voltage and slope factors are also quite variable across preparations (Figure 2). It should be noted that the magnitude of K^+^ conductances correlate well with corresponding channel gene mRNA copy numbers (Schulz et al., 2006), which serves as independent confirmation that variability is not solely due to noise or experimental error. We cannot provide a similar independent confirmation for variability in voltage-dependence, and it is not obvious to which degree half-activation and slope factor measurements may be more affected by experimental error than *g*_max_ is. However, variability in voltage-dependence may be due to post-translational modifications of ion channels (Jindal et al., 2008; Voolstra and Huber, 2014; Laedermann et al., 2015) or their phosphorylation state (Ismailov and Benos, 1995; Hofmann et al., 2014).

Not only did we not find any correlations between intrinsic and synaptic currents in the LP neuron, but we also did not find any correlations between intrinsic current parameters with either cycle period (Figure 5) or *φ*_LP on_ (Figure 6). This does not mean that intrinsic currents do not play an important role in controlling phase. Intrinsic properties, as determined by voltage-gated ionic currents, pump currents and even leak currents, are primary determinants of its activity. In the LP neuron, intrinsic properties have a great influence on *φ*_LP on_, as can be seen for example from the slow response of this neuron to repetitive dynamic clamp application of the same artificial synaptic input (Figure 7A). This slow response is indicative of a form of short-term memory over a timescale of many cycles that is attributed to intrinsic properties (Goaillard et al., 2010; Schneider et al., 2021). Some aspects of phase regulation are in fact dominated by intrinsic properties. For example, the phase difference between the LP and PY neurons is largely determined by differences in intrinsic currents, as experimentally applying identical synaptic input into both neuron types preserves their relative timing (Rabbah and Nadim, 2005).

Varying synaptic current amplitudes across individuals can still give rise to similar CPG output, for example in the leech heartbeat system (Norris et al., 2007; Norris et al., 2011). In the pyloric circuit, similar values for *φ*_LP on_ are achieved across individuals despite large variability of pacemaker synaptic input during ongoing rhythmic activity (Goaillard et al., 2009). We confirm the substantial variability in pacemaker to LP synaptic current amplitudes and in addition describe similar variability for PY to LP input (Figure 3C1). However, phase also depends on the relative timing, duration, and precise temporal trajectory of synaptic inputs (Prinz et al., 2003; Martinez et al., 2019b). In particular, *φ*_LP on_ is exquisitely sensitive to the shape and amplitude of synaptic input within preparations (Martinez et al., 2019b), and we show here that attributes describing the trajectory of the total synaptic current input to LP vary substantially across individuals (Figure 3C2-4). Therefore, similar values of *φ*_LP on_ are found across individuals despite varying intrinsic and synaptic currents.

In general, phase is dependent on an interplay of intrinsic and synaptic currents. Because *φ*_LP on_ adjusts over several cycles, any such interplay must occur at a much slower timescale than that of an individual cycle. Synaptic inhibition activates *I*_H_ and de-inactivates *I*_A_, which plays a critical role in determining rebound delay in follower neurons at different cycle periods (Harris-Warrick et al., 1995b; Harris-Warrick et al., 1995a; MacLean et al., 2005). *I*_H_ and *I*_A_ promote phase maintenance in individuals, particularly in conjunction with short-term synaptic depression, which results in an increase of inhibition with increasing cycle periods (Nadim and Manor, 2000; Manor et al., 2003; Bose et al., 2004; Greenberg and Manor, 2005; Mouser et al., 2008). Goaillard et al. (2009) recorded pyloric circuit activity and subsequently measured mRNA expression levels of the channel genes coding for *I*_H_ and *I*_A_ in LP, and also found no correlations with *φ*_LP on_. However, they did find *φ*_LP on_ to be correlated with the maximum value of a neuropeptide-activated current, which was also correlated with synaptic currents. Therefore, a lack of correlations between cycle period or *φ*_LP on_ and single intrinsic current parameters across individuals may simply mean that variability is well compensated across different currents.

The total synaptic current to the LP neuron is a combination of inputs from the pacemaker neurons AB and PD, and the 3-5 PY neurons, and therefore has a complex waveform shape (Figure 3). Of the 5 parameters that defined the synaptic waveform, three showed significant correlation with *P* across different animals (Figure 5), and four parameters had a strong correlation with *φ*_LP on_ (Figure 6). Surprisingly, these parameters did not include the strength of the synaptic input from the pacemaker or PY neurons. Goaillard et al. (2009) did not consider PY synaptic inputs to LP but separated AB and PD inputs by their different reversal potentials and found *φ*_LP on_ correlated with peak values of both, albeit with different sign. It is unclear whether this different finding simply results from the different way we defined synaptic strengths. However, the duty cycle and peak phase of the synapse, which strongly influence the LP phase in individuals (Martinez et al., 2019b) were among the correlated parameters. A linear dimensionality reduction using principal component analysis showed only two parameters (the first two principal components PC1 and PC2) sufficiently explained the correlation between cycle period and *φ*_LP on_. Consistent with these correlations, using the first two principal components to connect cycle period across preparations with the variability of *φ*_LP on_ was sufficient to explain phase maintenance across animals.

### Variability and co-regulation

Variability of intrinsic currents in STG neurons may be compensated by cell-type-specific co-regulation of different voltage-gated channels (Khorkova and Golowasch, 2007; Schulz et al., 2007; Temporal et al., 2012; Tran et al., 2019), but it is not known to which degree synaptic currents may be co-regulated. Variability of synaptic currents could be compensated for by variability in intrinsic currents, as has been suggested for the leech heartbeat system (Gunay et al., 2019), and as is implicit in theoretical work that shows similar circuit activity with different combinations of intrinsic and synaptic current levels (Prinz et al., 2004a; Onasch and Gjorgjieva, 2020). Alternatively, compensatory co-regulation of intrinsic currents could lead to consistent neuronal excitability on its own, and variability of synaptic trajectory then must be constrained to allow for consistent phases.

We found no evidence of co-regulation between intrinsic and synaptic currents (Figure 4), which suggests that phase constancy across preparations is not due to any obvious linear correlations that matched synaptic inputs to intrinsic properties. However, there are caveats to this analysis. We only performed pairwise linear correlations, and it is possible that we missed higher dimensional or nonlinear interactions. In addition, the nature of the intrinsic and synaptic current attributes we considered are somewhat mismatched. We described intrinsic voltage-gated currents with standard biophysical parameters, obtaining values for *g*_max_ and voltage-dependence. These parameters can be direct targets of cellular regulation, but it is not trivial to determine how their variability translates to variability in current magnitude and trajectory during ongoing circuit activity. In contrast, we assessed the magnitude and temporal trajectory of synaptic currents during ongoing pyloric activity, which are determined by pre- and postsynaptic properties as well as the voltage trajectories of the presynaptic neurons (Goaillard et al., 2009). Our synaptic current attributes therefore describe well the dynamics of synaptic interactions during circuit activity but can only serve as an indirect assessment of biophysical parameters that would be the targets of cellular regulation. In STG neurons, maximal synaptic currents or conductances and the dependence on presynaptic voltage have been assessed for their sensitivity to different neuromodulators (Zhao et al., 2011; Garcia et al., 2015; Li et al., 2018), but cannot be measured during ongoing circuit activity and their inter-individual variability has not been directly addressed.

### Correlation versus causation

Correlational analyses from spontaneous rhythmic activity restricted us to the limited variability of circuit output and did not afford us control of the variability of synaptic attributes. We therefore used the dynamic clamp, a technique that allows precise manipulation of synaptic inputs to individual neurons, which can be used to explore the role of a synapse in circuit activity (Bartos et al., 1999; Wright and Calabrese, 2011a, b; Martinez et al., 2019b). These experiments clearly showed that the LP burst onset is quite sensitive to the shape of the synaptic input waveform in a manner that was consistent across preparations. To our surprise, changing the waveform along PC1 or PC2, the two major directions of variability in the parameter space obtained from spontaneous activity, did not produce the largest influence on the LP burst onset. Instead, changing the waveform along PC5 and PC3, directions that did not show significant change with either cycle period or the LP burst onset across preparations, had the largest effect on the LP burst onset. In fact, when the waveform shape was kept constant along PC3, changing it along any other PC did not influence the LP burst onset at all (Figure 9D). Conversely, changing the waveform along PC3, while keeping any other PC constant, produced the strongest effects on the LP burst onset. This clearly indicated that, across preparations, the synaptic waveform was tuned by the circuit to remain unchanged along this direction of maximum sensitivity. Thus, in this sub-circuit, phase constancy across preparations is achieved partly by a precise control of the synaptic parameters that have the largest influence on phase.

It is tempting to interpret the correlation of underlying properties with attributes of circuit output as an indication that these attributes are controlled by these properties. However, properties may simply change with circuit output and not determine it. Similarly, one may interpret the lack of correlation as a sign of absence of influence. However, functional influence may be masked by the necessity to simply constrain parameters with large influence on output to a range that keeps output stable. The correlational relationships between the synaptic parameters and cycle period or *φ*_LP on_ that we described from spontaneous circuit output could statistically explain phase maintenance across animals. However, this explanation does not hold the test of causation. The same parameters that statistically predict *φ*_LP on_, or vary systematically with cycle period, have little influence on the burst onset when varied experimentally. Conversely, we only found the synaptic waveform attributes that are important for the control of phase by systematically varying them experimentally. Thus, our correlational explanation (Figure 7E) is in fact a consistency argument: if some synaptic parameters change with cycle period, then the same parameter must also change with *φ*_LP on_ in a manner that predicts phase constancy.

Many neural processes are found to co-vary across animals and correlations are often argued to be essential for the function of neural circuits (Golowasch, 2019; Santin and Schulz, 2019). It is important to remember that, despite the levels of degeneracy observed in the parameter space defining circuit output (Goldman et al., 2001; Bucher et al., 2005b; Swensen and Bean, 2005), correlations may simply be coincidental to the fact that the varying parameters do not have a meaningful influence on the function of interest (Hudson and Prinz, 2010; O’Leary et al., 2013).

Considering the numerous parameters that can influence the output of a neural circuit, inter-individual variability is neither surprising nor avoidable. Yet a consistent output pattern requires some essential combination of circuit parameters to be tightly constrained. Those that are not show variability across individuals and, because of the constraints of the output pattern, are forced to co-vary with output quantities that may also be relatively unconstrained, such as cycle frequency. Thus, parameters correlated with circuit output may contribute little to the output pattern, but rather become correlated because of constraints on this pattern.

### Differential control of activity phases

We addressed here how activity phases can stay consistent under control conditions, i.e., in the same circuit state. However, synaptic function and activity phases can be different between different circuit states, for example through the influence of neuromodulators (Harris-Warrick, 2011; Marder, 2012; Bucher and Marder, 2013; Marder et al., 2014b; Nadim and Bucher, 2014; Daur et al., 2016; Brzosko et al., 2019). In motor systems, the functional impact of such adjustments can be particularly transparent, as circuit reconfiguration through neuromodulation is for example a core mechanism for adjusting locomotion gait and speed (Harris-Warrick, 2011; Miles and Sillar, 2011; Bucher et al., 2015; Kiehn, 2016; Grillner and El Manira, 2020). Neuromodulators can affect neurotransmitter release, receptor properties, and postsynaptic intrinsic response properties (Nadim and Bucher, 2014). In addition, synaptic function can change because the activity profile of the presynaptic neuron is modified, as has been shown for STG neurons (Johnson et al., 2005; Johnson et al., 2011; Zhao et al., 2011). All these actions of neuromodulators can alter the temporal trajectory of synaptic responses. Therefore, our results provide a useful framework for understanding which aspects of the temporal dynamics of synaptic inputs can be altered by neuromodulators to change phase, and which changes phase relationships can be robust to.

## Methods

### Experimental preparation

Adult male crabs (*Cancer borealis*) were acquired from local distributors and maintained in aquaria filled with chilled (12-13°C) artificial sea water until use. Crabs were anesthetized before dissection by placing them in ice for at least 20 minutes. The stomatogastric nervous system including the stomatogastric ganglion (STG), esophageal ganglion, the pair of commissural ganglia, and the motor nerves were dissected from the stomach and pinned to a saline filled, Sylgard-coated (Dow Corning) Petri dish (schematic in Figure 1A). The STG was desheathed, exposing the somata of the neurons for intracellular impalement. Preparations were superfused with chilled (10-13°C) physiological saline containing: 11 mM KCl, 440 mM NaCl, 13 mM CaCl_2_ · 2H_2_O, 26 mM MgCl_2_ · 6H_2_O, 11.2 mM Tris base, 5.1 mM maleic acid with a pH of 7.4.

### Extracellular recordings of rhythmic patterns

Extracellular recordings from identified motor nerves were performed using pairs of stainless steel electrodes, placed inside and outside of a petroleum jelly well created to electrically isolate a small section of the nerve, and amplified using a differential AC amplifier (A-M Systems, model 1700). All traces were digitized using a Digidata 1332 data acquisition board and recorded in pClamp 10 software (both Molecular Devices).

The activity of three neuron types was used to identify the triphasic pyloric pattern (Marder and Bucher, 2007). The two pyloric dilator (PD) neurons belong to the pyloric pacemaker group of neurons, and we therefore used their burst onset as the reference time that defined each cycle of activity. The pyloric constrictor neurons include the single lateral pyloric (LP) neuron and multiple pyloric (PY) neurons. The constrictor neurons are follower neurons that receive strong inhibition from the pacemaker group and rebound from this inhibition to produce bursting activity at different phases. Spontaneous rhythmic pyloric activity was recorded from the lateral ventricular nerve (*lvn*), the pyloric dilator nerve (*pdn*), and occasionally also from the pyloric nerve (*pyn*) (Fig. 1A, nomenclature after Maynard and Dando, 1974). The *lvn* contains the axons of all three neurons types, with LP action potentials easily identifiable by their large amplitude. The *pdn* contains only the axons of the PD neurons, and the *pyn* only those of the PY neurons.

### Intracellular recordings and voltage clamp

For Intracellular impalement of the LP neuron soma, glass microelectrodes were prepared using the Flaming-Brown micropipette puller (P97; Sutter Instruments) and filled with 0.6 M K_2_SO_4_ and 20 mM KCl, yielding electrode resistances of 10-30 MΩ. Individual pyloric neurons were sequentially impaled, and the LP neuron was identified by its activity pattern and correspondence of action potentials between the soma recording and the extracellular recording of the *lvn* (Figure 1A). Recordings were amplified using Axoclamp 2B and 900A amplifiers (Molecular Devices) and recorded alongside the extracellular signals in pClamp. For current measurements, the LP soma was simultaneously impaled with two electrodes, and membrane potential was controlled in two electrode voltage clamp mode.

### Measurements of voltage-gated currents

In LP and other pyloric neurons, three intrinsic voltage-gated currents are relatively straightforward to measure in the intact circuit, without pharmacological manipulation (Zhao and Golowasch, 2012): the high-threshold K^+^ current (*I*_HTK_), the fast transient K^+^ current (*I*_A_), and the hyperpolarization-activated inward current (*I*_H_).

*I*_HTK_, consisting of the delayed rectifier and calcium-dependent K^+^ currents (Khorkova and Golowasch, 2007), was measured from the responses to voltage steps following a ~270 ms pre-step to −40 mV to inactivate *I*_A_. Voltage steps (750 ms) were delivered from −60 mV to +30 mV, in increments of 10 mV. In addition to subtracting the baseline current at −40 mV, the current recorded from the smallest voltage step was used to estimate the leak current, scaled proportionally for all voltage steps, and subtracted offline. The persistent component (*I*_HTKp_) was measured by taking an average of current recorded during the last 70 ms of a voltage step (90-99% of step duration). The transient component (*I*_HTKt_) was measured by taking the current peak, recorded during the first 150 ms of the voltage step.

*I*_A_ was obtained by recording the total K^+^ current (*I*_Ktot_) and digitally subtracting the previously measured *I*_HTK_. The neuron was held at −80 mV to remove inactivation. *I*_Ktot_ was then activated using voltage steps from −60 mV to +40mV in 10 mV increments. After subtracting *I*_HTK_ from *I*_Ktot_, the difference current was baseline subtracted. Because these currents were recorded without blocking sodium currents, effects of spikes generated in the electrotonically relatively distant axon were seen in the I_A_ traces (Fig. 2A, see also Zhao and Golowasch, 2012). Before measuring the peak amplitude of the currents, we used a robust smoothing function to remove the action potential-mediated transients. The amplitude of *I*_A_ was measured as the maximum during the first 150 ms of the voltage step.

*I*_HTKp_, *I*_HTKt_, and *I*_A_ were converted into conductances using the voltage-current relationships and an estimated K^+^ reversal potential (*E*_K_) of −85 mV. We then fit a standard sigmoid equation to a plot of conductance over membrane potential:

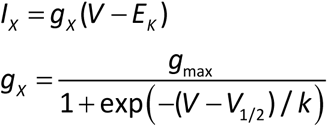

(X = HTKp, HTKt, or A). The sigmoid fits yielded values for maximal conductance (*g_max_*), voltage of half-activation (*V_1/2_*) and slope factor (*k*).

*I*_H_ was measured by holding LP at −40 mV for > 1.5 s and then stepping to more negative potentials between −60 mV and −120 mV for 5 s, in increments of 10 mV. Because of the small and variable size of *I*_H_ in the LP neuron, it is difficult to measure an accurate activation curve or reversal potential at physiological temperatures, particularly because rhythmic synaptic currents occur at similar amplitudes. Therefore, we only used the response to the step to −120 mV to estimate *I*_H_. The current was calculated by taking the difference between the current at the beginning and just before the end of the voltage step. The measured current was converted into conductance using a reversal potential of −30 mV (Buchholtz et al., 1992). In two preparations, the LP neuron did not have any measurable I_H_.

### Measurements of synaptic currents

Pyloric neurons receive mainly graded inhibitory synaptic input. Because LP is a follower neuron, pyloric oscillations continue while the LP neuron is voltage clamped, thus allowing for measurement of the IPSCs (Martinez et al., 2019b). LP was voltage clamped at a holding potential of −50 mV for at least 30 s. The current was averaged from the last 5 cycles measured, and a resulting unitary waveform was extracted. This unitary waveform was tagged at five distinct points, *t*_0_ to *t*_4_ (with the cycle period *P = t*_4_ − *t*_0_), which were connected using a piecewise linear graph (Figure 3B). The IPSC can be defined as the duration of this waveform from *t*_1_ to *t*_4_. The baseline of the IPSC (*I* = 0) was defined as the IPSC onset value at time *t*_1_. The IPSC waveform was normalized by *P*. Thus, the IPSC waveform can be characterized fully using the following parameters:

- Phase parameters:
  1. *DC*_LP_: duty cycle of the LP burst preceding the phases of synaptic input (= (*t*_1_ − *t*_0_) / *P*),
  2. *DC*_PY_: duty cycle of the PY component of the IPSC (= (*t*_2_ − *t*_1_) / *P*),
  3. *DC*_PD_: duty cycle of the pacemaker component of the IPSC (= (*t*_4_ − *t*_2_) / *P*),
  4. *θ*_LP_: peak phase of the synapse within the cycle, relative to the onset of the LP burst (= (*t*_3_ − *t*_0_) / *P*),
  5. *θ*_PD_: peak phase of the synapse within the cycle, relative to the onset of the PD burst (= (*t*_3_ − *t*_2_) / *P*),
  6. *Δ*_pk_: peak phase of the synapse within the IPSC (= (*t*_3_ − *t*_1_) / (*t*_4_ − *t*_1_)).
- Amplitude parameters:
  7. *I*_tot_: the maximum IPSC amplitude,
  8. *I*_PD_: amplitude of the pacemaker component of the IPSC,
  9. *I*_PY_: amplitude of the PY component of the IPSC (= *I*_tot_ - *I*_PD_).
- Slope parameters:
  10. *m*_PY_: rise slope of the PY component (= *I*_PY_ / (*t*_2_ - *t*_1_)),
  11. *m*_PD_: rise slope of the pacemaker component (= *I*_PD_ / (*t*_3_ − *t*_2_)),
  12. *m*_fall_: decay rate of the IPSC (= *I*_tot_ / (*t*_4_ − *t*_3_)).

Clearly, these parameters are not independent and include redundant ones. We defined all parameters in order to maintain the clarity of the biophysical interpretation of the IPSC and the contributing network components. However, for correlations between synaptic parameters and between synaptic and intrinsic current parameters, we defined the non-redundant subset, which consists of the following 5 parameters:

*DC*_PY_, *DC*_PD_, *Δ*_pk_, *m*_PY_, and *m*_fall_.

The other 7 parameters can be calculated from these values using simple geometry:

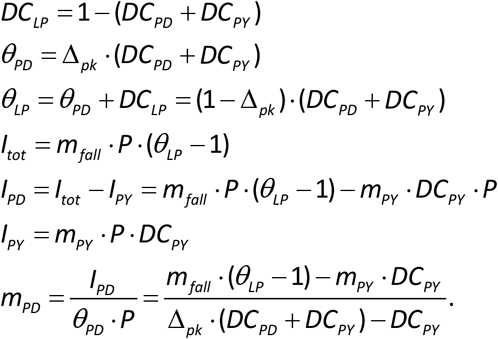

Note that the synaptic conductance waveform was taken to be identical to the synaptic current waveform measured in voltage clamp, as synaptic current at a constant holding potential simply scales with synaptic conductance.

### Dynamic clamp application of artificial synaptic input current

Dynamic clamp was implemented using the NetClamp software (Gotham Scientific) on a 64-bit Windows 7 PC using an NI PCI-6070-E board (National Instruments). We used dynamic clamp to inject artificial synaptic currents (*I*_syn_) into the synaptically isolated LP neuron (Prinz et al., 2004b; Zhao et al., 2010; Chen et al., 2016; Golowasch et al., 2017; Martinez et al., 2019b). In these experiments, the preparations were superfused with saline containing 10^−5^ M picrotoxin (Sigma Aldrich) to block the bulk of synaptic input to the LP neuron (Martinez et al., 2019a).

The dynamic clamp injected current *I*_syn_ was defined as

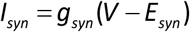

where *g*_syn_ is the synaptic conductance and *E*_syn_ is the synaptic reversal potential (set to −80 mV). *g*_syn_ was defined as a unitary stereotypical piecewise-linear waveform, mimicking the experimentally measured synaptic conductance. The unitary synaptic conductance waveforms were constructed using the following algorithm:

- *I*_tot_ = 1.
- *DC*_LP_ + *DC*_PY_ + *DC*_PD_ = 1.
- *DC*_LP_, *DC*_PY_, θ_PD_ and *I*_PY_ were chosen from the values between 0 and 1, in increments of 0.2.
- *m*_PD_ > *m*_PY_
- *DC*_LP_ + *DC*_PY_ + θ_PD_ < 1.

These rules yielded a total of 80 waveforms, including a few duplicates. Each waveform was applied periodically with a cycle period of 1 s and a peak amplitude of 0.4 *μ*S. In each trial, the artificial synaptic input was applied for at least 30 s.

In these experiments, the bursting activity of the LP neuron was quantified by measuring the latency of the burst onset compared to the end of the conductance waveform (*t*_4_ in Figure 3B1; latency shown in Figure 8A). Note that this is different from the burst latency measured for calculating the LP phase during an ongoing pyloric rhythm (Figure 1A, right panel), which is measured with respect to the onset of the pacemaker PD neuron bursts. However, our primary goal in these experiments was to understand how changing the shape of the synaptic input influenced the activity of the LP neuron. The corresponding reference point in the dynamic clamp experiments would have been the onset of the pacemaker component of the synaptic input (*t*_2_ in Figure 3B1). However, had we measured latency with respect to *t*_2_, our calculation of latency would have given the appearance that it changes with the waveforms, even if there was absolutely no change in the LP neuron activity. This is because *t*_2_ is quite different across the 80 waveforms (Figure 8B). The end of the conductance waveform is the only reference point that accurately reports changes in the LP activity due to the waveform shape is the end timepoint of the synaptic waveform.

## Data analysis

All analysis was performed using custom scripts written in MATLAB (MathWorks). All linear correlations were measured using MATLAB built-in function ‘corr’, which computes Pearson’s linear correlation coefficient. Principal component analysis was performed using the MATLAB ‘pca’ function. Figures were plotted in MATLAB and panels were assembled in CorelDRAW (version 2020, Corel).

The activity phase (*φ*_LP on_) of the LP neuron burst onset is defined as the time interval between the onset of the pacemaker PD neuron’s burst to the onset of the LP neuron burst, normalized by the period (*P*) of that cycle, defined as the time interval between the two consequent PD neuron bursts (Figure. 1A). To examine the effect of changing the synaptic waveform along each principal component (using dynamic clamp) on *φ*_LP on_, for each principal component PC_j_ (j = 1,…,5), we projected all 80 synaptic waveforms onto the plane defined by PC_j_ and each PC_k_ (k ≠ j). We then found all waveform pairs (say *w_n_* and *w_m_*) that fell within ±0.1 of each PC_k_ value and were different by at least 0.1 in PC_j_, and measured *φ*_LP on_ for each waveform. (In this analysis, to exclude any effect of the duration of inhibition, we computed *φ*_LP on_ by calculating the latency as the time-to-first-spike of the burst relative to the end of dynamic clamp inhibition, and then divided this latency by *P*.) We then calculated the sensitivity of the LP burst onset latency (lat) for this pair of waveforms *w_n_* and *w_m_* as

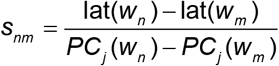

(here, PC_j_(*w_n_*) is assumed to be > PC_j_(*w_m_*)). We reported the sensitivity of *φ*_LP on_ to PC_j_, while keeping PC_k_ constant, as the statistical distribution defined by *s_nm_* values in all preparations (200-500 data points, depending on j and k). The overall sensitivity burst latency to PC_j_ in each preparation was calculated as the mean value of all *s_nm_* values when changing PC_j_, while keeping PC_k_ constant, for all k ≠ j, in that preparation.

## Acknowledgements

This study was supported by NIH MH060605.

---

**Figure 2—figure supplement 1.**
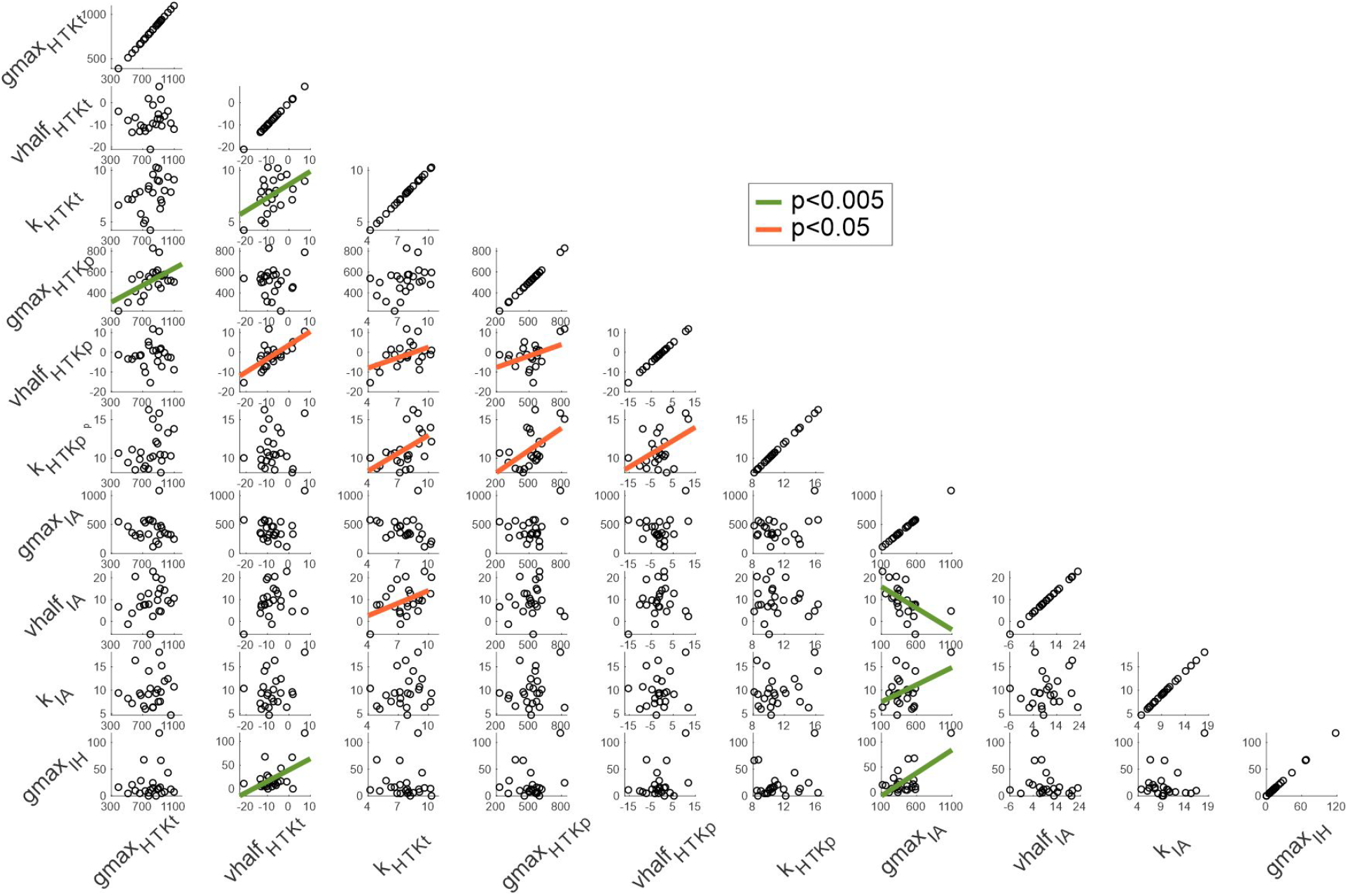
Correlations among parameters of the voltage-gated ionic currents. No fit line in a panel indicates lack of correlation.

